# Development and Refinement of Microbial DNA Extraction Protocol from Bovine Milk

**DOI:** 10.64898/2026.04.30.721090

**Authors:** Rowan Cook, Joana Lima, Richard J. Dewhurst, Sharon A. Huws, Christopher J. Creevey, Holly J. Ferguson

## Abstract

Milk is a challenging matrix to extract sufficient microbial DNA from for downstream analysis. This study assessed fourteen DNA extraction protocols for their DNA outputs. An adaptation of the QIAGEN DNeasy PowerSoil kit, which increased initial sample volume and maintained all volume of lysate following bead beating proved most effective.

## Introduction

The development of next-generation sequencing techniques has facilitated the characterisation of the bovine mammary microbiome, contradicting previously held beliefs that the udder was sterile unless diseased (Ruegg, 2022; Reuben and Torres, 2025). Microbiome sequencing has generated novel insights into areas such as mastitis health and colostrum quality, however, obtaining sufficient DNA extracts from milk has proved challenging (Pirondini *et al*., 2010; Oikonomou *et al*., 2014; Falentin *et al*., 2016; Pollock *et al*., 2021; Schwenker *et al*., 2022; Scully *et al*., 2025). Schwenker *et al*. (2022) suggested possible reasons for this could be the physical, chemical, and biological characteristics of the milk matrix including the composition of fat, protein, and calcium which inhibit enzymatic and PCR processes.

Different methods for DNA extraction from milk have previously been explored, with an array of detergents, chemical, and mechanical lysis evaluated (Cremonesi *et al*., 2006; Quigley *et al*., 2012; Volk *et al*., 2014; Schwenker *et al*., 2022). The difference between fractional compositions of the milk have also been explored for DNA extraction yields and microbial sequencing results (Lima *et al*., 2018). Considering the fractional composition is important, as targeting the pellet during extraction may bias towards identification of pathogenic bacteria located within the pellet matrix, or targeting the cream-layer may over-inflate numbers of Staphylococci bacteria, reported to bind to fat globules (Ali-Vehmas *et al*., 1997; Hickey *et al*., 2015).

Commercial kits have been developed which claim to be suitable for milk DNA extraction as an alternative to published research protocols using phenol-chloroform extraction with alcohol precipitation (Murphy *et al*., 2002; Volk *et al*., 2014; Schwenker *et al*., 2022). This provides the convenience of a ready-made kit with consumables included and reduces exposure to harmful chemicals (Quigley *et al*., 2012).

The development of a robust DNA extraction method is required to enable reliable characterisation of the milk microbiome to enable the investigation of its potential to improve animal health and productivity whilst reducing antimicrobial usage. Therefore, this study compared fourteen different DNA extraction protocols to determine the optimal approach for milk samples.

## Materials and Methods

Milk samples were obtained from Holstein-Friesian dairy cattle at Scotland’s Rural College’s (SRUC) Dairy Research and Innovation Centre (Dumfries, United Kingdom). Samples were collected aseptically following the National Mastitis Council (2004) guidelines. Animals were selected based on mastitis health status; diagnosed with clinical mastitis signs (n = 3), an elevated somatic cell count (SCC) > 200,000 cells/ml indicating subclinical mastitis (n = 4), or healthy (no clinical mastitis signs and SCC < 200,000 cells/ml, n = 4). Animals were milk recorded every two weeks by National Milk Records and Cattle Information Services, alternately.

All fourteen methods were derived from the Newton protocol (Newton, 2023, supplementary 1), the QIAGEN Stool Kit (pathogen protocol, cat. no. 51604, QIAGEN, Hilden, Germany), or the QIAGEN Soil Kit (cat. no. 47014), and were refined through a structured, stepwise optimisation process, with methods that did not produce viable outcomes being excluded from subsequent testing. Whole milk was used for testing unless a preprocessing step was specifically described which targeted a specific constituent. A description of the DNA extraction protocols can be seen in Table 1. Protocols 1-9 were tested on clinical mastitis, subclinical mastitis and healthy samples, and protocols 10-14 tested only on subclinical and healthy samples as these proved to be particularly challenging, and were therefore the focus of the optimisation.

**Table 1.** Microbial DNA extraction protocols assessed.

| **Protocol** | **Base of protocol** | **N** | **Adaptions** |
| --- | --- | --- | --- |
| **1** | Newton (2023), Supplementary 1 | 15 | Centrifugation speeds were adjusted to accommodate available laboratory equipment (9,500 x g maximum). |
| **2** | QIAGEN QIAamp Fast DNA Stool kit | 6 | No adaptions from kit protocol |
| **3** | QIAGEN QIAamp Fast DNA Stool kit | 6 | Preprocessing: 5 ml of milk was centrifuged and the fat layer removed, with the centrifugation and fat removal repeated once more. Supernatant was removed and 1 ml of PBS added to pellet before being vortexed and transferred to a mct for centrifugation (9,500 x g, two minutes). The supernatant was then used following the manufacturers pathogen protocol. |
| **4** | Newton (2023) | 4 | Increased volume of sample from 100 to 500 µl. |
| **5** | Newton (2023) | 4 | QIAGEN Powerbead Pro tube was used to bead beat sample and inhibitEX buffer mixture for three minutes. |
| **6** | Newton (2023) | 3 | Combination of protocols 4 and 5. |
| **7** | Newton (2023) | 4 | Increased sample volume to 500 µl and bead beat without inhibitEX buffer. |
| **8** | Newton (2023) | 4 | Identical to protocol 5 but replaced inhibitEX buffer with CD1 (QIAGEN PowerSoil pro kit) |
| **9** | Newton (2023) | 4 | Sample volume was increased to 500 µl. In the previous protocol, combining 600 µl of supernatant with proteinase K resulted in approximately 600 µl of supernatant remaining, which was discarded. In this protocol, the entire 1200 µl of supernatant was retained and carried forward in two microcentrifuge tubes. All lysate was subsequently processed through a single spin column per sample. |
| 10 | Newton (2023) | 2 | Increased sample volume to 500 µl and vortexed for 20 minutes before mixing with inhibitEX. |
| 11 | Newton (2023) | 2 | 2 ml of milk sample was centrifuged for ten minutes (9,500 x g) and the supernatant transferred to a new mct. The fat and pellet were resuspended with 1 ml PBS, centrifuged as before and the supernatant added to the first supernatant. Protocol was then carried out with 500 µl of the combined supernatant |
| 12 | Newton (2023) | 2 | Combination of protocols 10 and 11. |
| 13 | QIAGEN DNeasy PowerSoil Pro kit | 2 | Increased starting volume to 500 µl. Approximately 1200 µl of supernatant was produced after bead beating which was split evenly between two clean mcts. Protocol continued per kit instructions but with two tubes per sample and combined all lysate per sample into the same spin column. |
| 14 | QIAGEN DNeasy PowerSoil Pro kit | 2 | Similar to protocol 13 but increased the starting sample volume to 1800 µl divided into two Powerbead Pro tubes. After bead beating, approximately 3000 µl of supernatant is produced which was aliquoted into five mcts (~600 µl each). Protocol continued with five tubes per sample and combined all lysate per sample into the same pin column. |

**N IS THE NUMBER OF SAMPLES TESTED. ALL BEAD BEATING WAS CONDUCTED USING QIAGEN POWERBEAD PRO TUBES ON A VORTEX AT MAXIMUM SPEED (3,200 RPM) FOR THREE MINUTES. MCT = MICROCENTRIFUGE TUBE**

DNA quantification was conducted using an Invitrogen Qubit 4 fluorometer (Thermo Fisher, Massachusetts, USA) using a High Sensitivity Qubit™ double-stranded (ds) DNA Quantification Assay Kit, following the kit protocol.

## Results

Clinical mastitis samples had the highest DNA yield across all extraction protocols (171.80 ± 213.03 ng/µl), followed by healthy samples (1.84 ± 2.19 ng/µl) and subclinical mastitis samples (0.80 ± 2.65 ng/µl). Protocol 6 produced the highest average DNA yield overall (248.12 ± 429.45 ng/µl), but protocol 14 produced the highest yield when considering healthy and subclinical mastitis samples only (7.84 ± 2.60 ng/µl, Figure 1).

**Figure 1.**
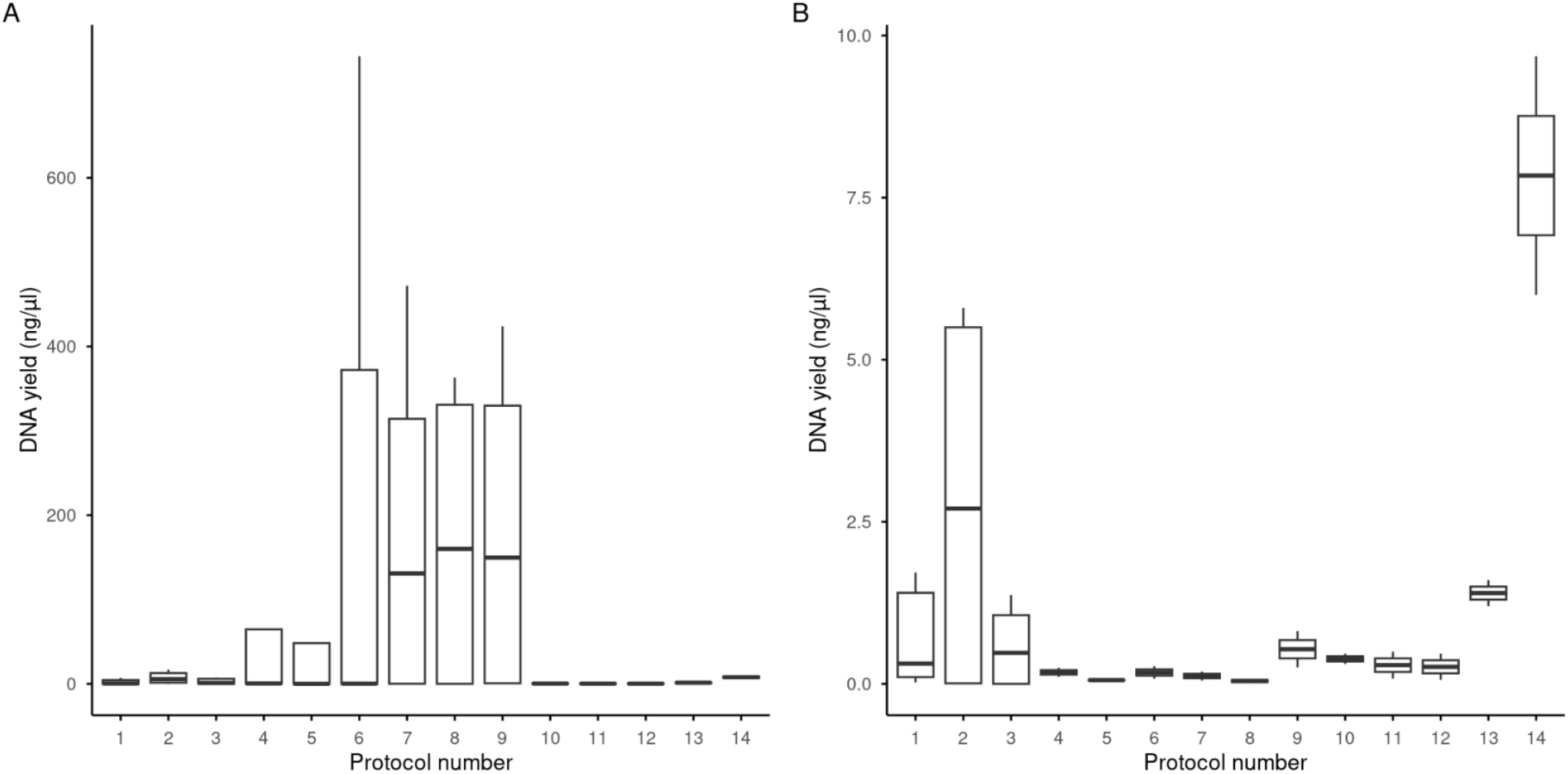
DNA extraction yields of the fourteen different protocols tested. Boxplots A and B show the DNA extraction yields including and excluding clinical mastitis samples, respectively.

## Discussion

Protocol 14 was the most effective method for extracting microbial DNA from healthy and subclinical mastitis milk samples. An illustrative schematic of this protocol is shown in Figure 2. In addition to improved performance, this protocol was less time-consuming than the Newton-based and Qiagen stool kit-based protocols. Following its development, protocol 14 was used for DNA extractions from 160 milk samples, including healthy, subclinical, and clinical mastitis milk, consistently yielding sufficient DNA for both shotgun metagenomic sequencing and amplicon sequencing (16S rRNA and ITS targets).

**Figure 2.**
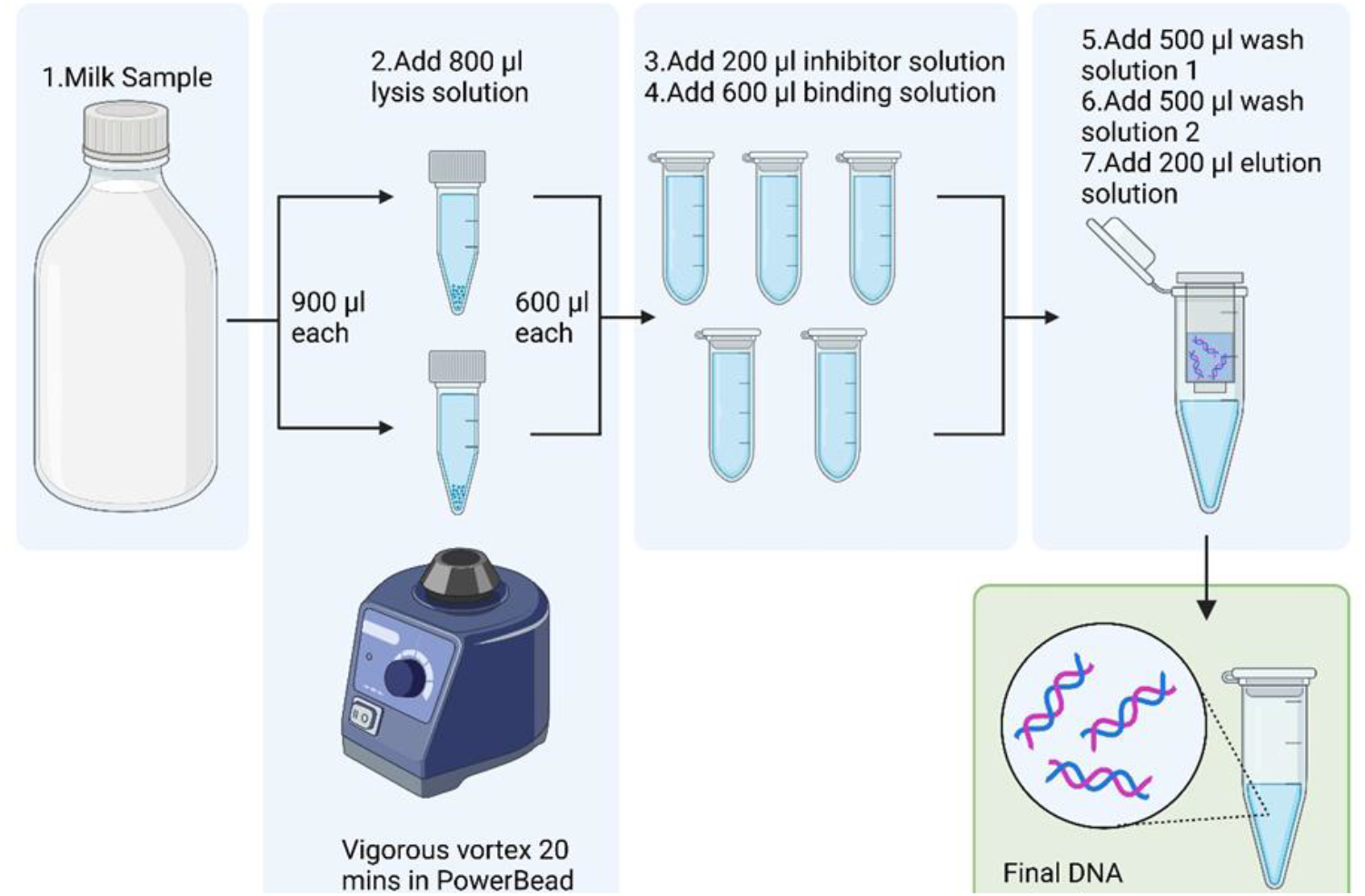
Milk DNA extraction protocol adapted from Qiagen DNeasy PowerSoil Pro kit. Created in BioRender. Cook, R. (2025) https://BioRender.com/t67o674.

The success of this protocol is likely attributable to the increased in initial sample volume, which provided a greater quantity of microbial DNA for extraction, and to the retention of the full lysate following bead-beating, minimising DNA loss during early processing steps.

Milk is a low-biomass matrix in which microbial DNA is present in considerably lower concentrations than host DNA (Pollock *et al*., 2021). Extraction efficiency is further compromised by the presence of fat, protein, and calcium ions that can inhibit downstream enzymatic reactions (Lima *et al*., 2018). Previous studies focused on the milk microbiome targeted either the milk pellet or milk pellet and fat layer together for DNA extraction and discard the skimmed milk to minimise the impact of these compromising components (Lima *et al*., 2018). We assessed these methods in protocols 3, 10, 11 but did not find them to be as effective as protocol 14 which targeted whole milk.

The performance protocol 14 across multiple sample types (i.e. healthy vs mastitis) and its successful use with multiple sequencing approaches supports its suitability for high-throughput milk microbiome studies. This method was not validated on other milk sources such as colostrum or bulk tank milk, therefore further optimisation may be required for these matrices.

## Conclusion

Of the fourteen protocols assessed in this study, the modified QIAGEN DNeasy PowerSoil Pro protocol, which increased the initial sample volume to 1800 µl and retained all lysate following bead beating, produced the highest DNA yields. The development of this method is particularly useful in the microbial characterisation of healthy and subclinical milk samples, from which extract sufficient DNA yield is challenging. The protocol utilises whole milk without the need for preprocessing prior to extraction.

## Data availability

The data that support the findings of this study are available from the corresponding author upon request.

## Acknowledgements

Authors wish to thank the farm and technical staff at SRUC’s Crichton Royal Farm for their help in sample collection. Additional thanks to E.E. Newton for kindly sharing his DNA extraction protocol.

## Author contributions: CRediT

**Rowan Cook**: Conceptualization; writing – original draft; investigation; formal analysis; writing – review and editing; Data curation. **Christopher J. Creevey:** Review and editing; supervision; Funding acquisition. **Richard J. Dewhurst:** Review and editing; supervision, Funding acquisition. **Holly J. Ferguson**: Funding acquisition; conceptualization; review and editing; supervision, Project administration. **Sharon A. Huws:** Review and editing; supervision; Funding acquisition. **Joana Lima:** conceptualization; review and editing; supervision; Data curation; Investigation.

## References

Ali-Vehmas, T., Westphalen, P., Myllys, V. and Sandholm, M. (1997). Binding of Staphylococcus aureus to milk fat globules increases resistance to penicillin-G. Journal of Dairy Research, 64, 253–260.

Cremonesi, P., Castiglioni, B., Malferrari, G., Biunno, I., Vimercati, C., Moroni, P., Morandi, S. and Luzzana, M. (2006). Improved method for rapid DNA extraction of mastitis pathogens directly from milk. Journal of Dairy Science, 89, 163–169.

Falentin, H., Rault, L., Nicolas, A., Bouchard, D.S., Lassalas, J., Lamberton, P., Aubry, J.-M., Marnet, P.-G., Le Loir, Y. and Even, S. (2016). Bovine teat microbiome analysis revealed reduced alpha diversity and significant changes in taxonomic profiles in quarters with a history of mastitis. Frontiers in Microbiology, 7, 480.

Hickey, C.D., Sheehan, J.J., Wilkinson, M.G. and Auty, M.A.E. (2015). Growth and location of bacterial colonies within dairy foods using microscopy techniques: a review. Frontiers in Microbiology, 6.

Lima, S.F., Bicalho, M.L.D.S. and Bicalho, R.C. (2018). Evaluation of milk sample fractions for characterization of milk microbiota from healthy and clinical mastitis cows.(ed JJ Loor) PLOS ONE, 13, e0193671.

Murphy, M.A., Shariflou, M.R. and Moran, C. (2002). High quality genomic DNA extraction from large milk samples. Journal of Dairy Research, 69, 645–649.

National Mastitis Council. (2004). Microbiological Procedures for the Diagnosis of Bovine Udder Infection and Determination of Milk Quality. National Mastitis Council, Madison, WI, USA.

Newton, E.E. (2023). The Effect of Phycological Supplementation of Dairy Cow Diets on Milk Quality in European Dairy Systems. PhD Thesis, University of Reading, Reading, United Kingdom.

Oikonomou, G., Bicalho, M.L., Meira, E., Rossi, R.E., Foditsch, C., Machado, V.S., Teixeira, A.G.V., Santisteban, C., Schukken, Y.H. and Bicalho, R.C. (2014). Microbiota of Cow’s Milk; Distinguishing Healthy, Sub-Clinically and Clinically Diseased Quarters.(ed LL Guan) PLoS ONE, 9, e85904.

Pirondini, A., Bonas, U., Maestri, E., Visioli, G., Marmiroli, M. and Marmiroli, N. (2010). Yield and amplificability of different DNA extraction procedures for traceability in the dairy food chain. Food Control, 21, 663–668.

Pollock, J., Salter, S.J., Nixon, R. and Hutchings, M.R. (2021). Milk microbiome in dairy cattle and the challenges of low microbial biomass and exogenous contamination. Animal Microbiome, 3.

Quigley, L., O’Sullivan, O., Beresford, T.P., Paul Ross, R., Fitzgerald, G.F. and Cotter, P.D. (2012). A comparison of methods used to extract bacterial DNA from raw milk and raw milk cheese: DNA extraction from raw milk and cheese. Journal of Applied Microbiology, 113, 96–105.

Reuben, R.C. and Torres, C. (2025). Integrating the milk microbiome signatures in mastitis: Milk-omics and functional implications. World Journal of Microbiology and Biotechnology, 41, 41.

Ruegg, P.L. (2022). The bovine milk microbiome–an evolving science. Domestic Animal Endocrinology, 79, 106708.

Schwenker, J.A., Friedrichsen, M., Waschina, S., Bang, C., Franke, A., Mayer, R. and Hölzel, C.S. (2022). Bovine milk microbiota: Evaluation of different DNA extraction protocols for challenging samples. MicrobiologyOpen, 11, e1275.

Scully, S., Earley, B., Smith, P.E., Finnie, M.S.J., McAloon, C., Buckley, F., Kenny, D.A. and Waters, S.M. (2025). Characterisation of the bacterial and archaeal microbiota in fresh colostrum collected from a single, spring-calving dairy herd.(ed MSV Salles) PLOS One, 20, e0335718.

Volk, H., Piskernik, S., Kurinčič, M., Klančnik, A., Toplak, N. and Jeršek, B. (2014). Evaluation of different methods for DNA extraction from milk. Journal of Food and Nutrition Research, 53, 97–104.

