## Supplementary 1 for "Development and Refinement of Microbial DNA Extraction Protocol from Bovine Milk"

This protocol follows a milk DNA extraction method proposed by E.E. Newton (University of Reading) and described in his PhD thesis (Newton, 2023). All centrifugation speeds were adjusted from 16,000 x g to 9,500 x g to accommodate available laboratory equipment.

Milk samples were thawed at room temperature. An aliquot of 100 µl was transferred to a 2 ml microcentrifuge tube containing 1 ml InhibitEX buffer (QIAGEN QIAamp Fast DNA Stool kit, Hilden, Germany). Samples were vortexed at 3200 rpm for 1 min and incubated in a water bath at 90 °C for 5 min. Following incubation, samples were vortexed for 15 s and centrifuged for 1 min to separate the fat layer. The fat layer was carefully removed, and the remaining supernatant transferred to a new 2 ml microcentrifuge tube.

RNase A (4 µl; Fisher Scientific, Loughborough, UK) was added to each sample and incubated for 3 min at room temperature to reduce RNA contamination. Samples were then centrifuged for 3 min, and 600 µl of the supernatant was transferred to a new 2 ml microcentrifuge tube containing 25 µl proteinase K (QIAGEN QIAamp Fast DNA Stool kit). Buffer AL (600 µl; QIAGEN QIAGEN QIAamp Fast DNA Stool kit) was added, and samples were incubated at 70 °C for 10 min. Ethanol (600 µl, 99%) was added, and samples were vortexed for 15 s.

The resulting lysate was applied to a QIAGEN spin column in 600 µl aliquots and centrifuged for 1 min per aliquot, discarding the flow-through between centrifugation steps. Wash steps were performed using 500 µl Buffer AW1 (QIAGEN QIAamp Fast DNA Stool kit), centrifuged for 1 min, followed by 500 µl Buffer AW2 (QIAGEN QIAamp Fast DNA Stool kit), centrifuged for 3 min.

DNA was eluted by adding 50 µl Buffer EB (QIAGEN, not included in kit) directly to the spin column membrane and incubating for 2 min at room temperature prior to centrifugation for 1 min. The eluate was reapplied to the spin column, incubated for an additional 1 min, and centrifuged again for 1 min to maximise DNA recovery.
